# Botanical nursing by a dwarf shrub as a nature-based approach to plant biodiversity conservation in the Nepal’s Himalayas

**DOI:** 10.64898/2025.12.23.696287

**Authors:** Rabindra Parajuli, Bishnu Timilsina, Rita Chhetri, Prakash Bhattarai, Khadananda Acharya, Charles B. van Rees

## Abstract

Plant-plant positive interactions are among the main drivers of plant community structure, especially in stressful environments. While the ecological dynamics of facilitation are increasingly understood, studies evaluating the importance of nurse plants in preserving phylogenetic diversity and protecting species of medicinal value and conservation concern are still limited. Here, we investigated the nurse effects of a Himalayan endemic shrub, *Berberis angulosa*, in promoting taxonomic richness, composition and phylogenetic diversity of overall and highly valued species in the high-elevation mountain landscape of Langtang National Park, Nepal. Using pair-wise sampling design, we recorded plant communities within the patches of *Berberis* shrubs and in open gaps at three altitudinal belts.

Compared to adjacent open areas, *Berberis* patches were found to harbor compositionally distinct and phylogenetically diverse plant communities with significantly higher average overall and human-valued species richness. Relative interaction indices showed clear patterns along elevation gradient with the strongest facilitation observed at highest elevation. Similarly, Net Relatedness Indices (NRI) were significantly different between nursed vs open habitats for both overall and human-valued species, with lower mean NRI within *Berberis* patches indicating higher phylogenetic diversity of communities within the nursed habitats. Facilitation by *Berberis* enhanced community-level richness by 40% (39 species; total = 97), including 19 human-valued species (∼36%; total = 53) that occurred exclusively within shrub canopies. Many medicinal, endemic and conservation priority species were either shrub specialists or had significant association with *Berberis* patches, illustrating the protective role of the nurse shrub. Our results demonstrate that the facilitative effect of *Berberis* is not only crucial in maintaining plant communities with diverse evolutionary histories but also equally contributing to local beta and landscape-scale gamma biodiversity, sustaining valuable ecosystem services and benefits to human communities in the Himalayas. These findings suggest that appropriate management of such nurse plants may constitute a nature-based approach to biodiversity conservation, that should be considered in policies for sustainability, resilience and ecosystem functioning in this and similar alpine systems.

## INTRODUCTION

Facilitation shapes community structure across environmental and geographic scales, affecting biological diversity across taxonomic, phylogenetic, and functional dimensions (Callaway 2007; Valiente-Banuet and Verdú 2007; Butterfield et al. 2013; Cavieres et al. 2014; Losapio et al. 2021; Bashirzadeh et al. 2022). Among plants, these interactions manifest such that one or more species, often called nurse plants, promote the fitness of other, less environmentally tolerant (i.e. stress-sensitive) species, mostly in severe environmental conditions (*sensu* Stress Gradient Hypothesis – SGH, (Bertness and Callaway 1994). Nurse plants ameliorate bio-physical stresses and create favorable micro-habitats beneath their canopies, facilitating the growth, survival and recruitment of other species, thus supporting the formation of unique phylogenetic and plant community structure (Callaway 2007; Valiente-Banuet and Verdú 2007; Butterfield et al. 2013; Kikvidze et al. 2015; Pistón et al. 2016).

Phylogenetic diversity (PD) quantifies the evolutionary relationships among species within a community and serves as a fundamental metric to understand biodiversity patterns from a historical perspective and their contributions to ecosystem stability and resilience (Faith 1992). Conserving PD is therefore essential to preserve the evolutionary history of an ecosystem, maintain adaptability to environmental change, support critical ecosystem services, and preserve units of evolutionary history (Srivastava et al. 2012). Facilitation by nurse plants is effective in maintaining phylogenetic diversity in stressful environments, including alpine mountains and degraded ecosystems, by creating microhabitats that support the coexistence of phylogenetically distinct species under harsh conditions (Butterfield et al. 2013; Navarro-Cano et al. 2016; Pistón et al. 2016). The extent of facilitation in conserving PD, however, varies across ecosystem types (Bashirzadeh et al. 2022), underscoring the need to extend research across a wider geographic regions.

The Himalayas, the youngest and highest mountain range on Earth (Roy and Purohit 2018), remain among the least explored systems, where the implications of plant facilitation on phylogenetic diversity in these extreme montane environments remain poorly studied. They harbor unique ecosystems and diverse habitats exhibiting high endemism and encompassing distinct evolutionary lineages, thereby contributing to remarkable phylogenetic diversity (Ahmad et al. 2025). However, the Himalayan mountains are vulnerable to biodiversity loss due to the ongoing climate change, habitat fragmentation and intensifying resource exploitation (Myers et al. 2000; Xu et al. 2009). It is therefore critical to safeguard this diversity against various global and local stressors (Ahmad et al. 2025), and facilitation by nurse shrubs may offer one of the nature-based conservation strategy to this end, while providing opportunities for better understanding this ecological phenomenon (Parajuli et al. 2021). Despite increasing knowledge on how facilitation shapes the phylogenetic diversity in other critical ecosystems (Valiente-Banuet and Verdú 2007; Butterfield et al. 2013; Navarro-Cano et al. 2016; Bashirzadeh et al. 2022), the Himalayan region remains a major knowledge gap in the geographically biased corpus of literature on this topic, necessitating research to better understand and safeguard its biological heritage (Ahmad et al. 2025).

Importantly, there is some debate and uncertainty regarding the prevalence of facilitative interactions and their importance in structuring the plant communities in the Himalayan region (e.g., (de Bello et al. 2011; Dvorský et al. 2013; Liancourt et al. 2017; Ram and Chawla 2024). For instance, (de Bello et al. 2011), (Dvorský et al. 2013) and (Liancourt et al. 2017) showed that competitive interactions, rather than facilitative ones, were more common in the environmentally severe western arid Himalayas and Karakoram in Ladakh, India, contradicting the SGH. Similarly, (Ram and Chawla 2024) found no evidence of facilitation by several dwarf shrubs (*Caragana versicolor*, *Juniperus polycarpos* and *Rhododendron anthopogon*) on herbaceous plant communities in alpine regions of the western Himalayas of India. However, (Iyengar et al. 2017) found that a dwarf compact shrub (*Caragana versicolor*) facilitated rare species and increased local herbaceous plant communities in the western trans-Himalayan region of Spiti, India. Likewise, in the central and eastern Himalayas (including the Hengdung mountains) and Tibetan Plateau, facilitation by cushion plants and shrubs strongly influenced plant community structure, with increasing interaction intensity observed where environmental stresses were higher (e.g., (Yang et al. 2010; Pugnaire et al. 2015; Chen et al. 2015; Ale et al. 2018, 2023; Parajuli et al. 2021).

Characterized by high cultural diversity (Chaudhary 1998; Xu et al. 2009; Negi et al. 2025), the Himalayan region hosts unique ethnoecological interactions due to local communities’ reliance on natural resources for socio-ecological resilience and sustainability (Chaudhary 1998; Ghimire et al. 2004, 2008a; Aryal et al. 2014; Negi et al. 2025). In Nepal, people depend heavily on plant resources such as medicinal herbs, and pastures used in transhumance – a traditional practice involving the seasonal movement of livestock between high- and low-altitude pastures to optimize grazing, as integral parts of their livelihood (Ghimire et al. 2004; Aryal et al. 2014). These landscapes are culturally shaped by anthropogenic disturbances, such as transhumance, which act as ecological drivers of plant diversity (Aryal et al. 2015). In this system, for example, both overgrazing and the complete absence of grazing can lead to declines in plant diversity (Sharma et al. 2014; Aryal et al. 2015). In addition to changing anthropogenic influences on the landscape, alpine ecosystems like the Himalayas are experiencing the most severe impacts of climate change and its cascading effects on biodiversity and ecosystem functioning (Xu et al. 2009; Shrestha et al. 2012; Bhattarai et al. 2022; Ahmad et al. 2025). Moreover, by altering the availability of resources, including medicinal and high-value plants, climate and land use changes may impact local livelihood and ultimately threaten socioeconomic sustainability and resilience (Xu et al. 2009; Aryal et al. 2014; Aryal 2015; Negi et al. 2025).

Facilitation may play an important role as a nature-based conservation measure in these imperiled Himalayan socio-ecological systems, where microhabitats created by nurse plants provide shelter for culturally or economically valuable herbs and species of conservation concern (Parajuli 2012; Kikvidze et al., 2015; Cavieres et al., 2016; Parajuli et al., 2021). Despite the potential for such studies, research on facilitation as a nature-based tool to enrich ecosystem services, sustain ethnoecological ties, contribute to local livelihoods, and maintain biodiversity across multiple dimensions remains limited. Limited research concerning the implications of facilitation in the phylogenetic diversity of plant communities, including high value plants, along with debate over its prevalence in this system, necessitate further exploration to better understand how facilitation influences plant communities and their evolutionary relationships in the Himalayas.

Here, we investigated the role of facilitation by a Himalayan endemic dwarf shrub, *Berberis angulosa* (hereafter, *Berberis*), on plant community composition, and taxonomic and phylogenetic diversities of both overall and human-valued species, including medicinal herbs, endemic species and conservation priority taxa. We predicted that facilitation by *Berberis* supports compositionally and phylogenetically distinct taxa, thus conserving plant biodiversity and ecosystem functioning in the human-adapted high-elevation alpine mountains. While testing our prediction, we specifically address the following questions:

a. What are the patterns of plant-plant interactions along elevation gradients in the study region?
b. Are there more high-value medicinal, endemic, and conservation priority species associated with Berberis patches than open gaps?
c. Does facilitation by native nurse shrubs contribute to conserving phylogenetic diversity in Nepal’s Himalaya?

## MATERIALS AND METHODS

### Study area

The study was conducted in the Gosaikunda sector, a Himalayan eco-cultural landscape within Langtang National Park (LNP) of Central Nepal (Fig. 1). LNP is one of the biodiverse areas of the country with rich species diversity and endemism, harboring a high number of rare, endemic and medicinal species (Ghimire et al. 2008b; Joshi and Joshi 2022). Of the 47 endemic plant species recorded in the park, 18 are found in the Gosaikunda sector only (Ghimire et al. 2008b), highlighting the critical importance of research encompassing various facets of biodiversity in the region. The unique geography, diverse topography, and rich assemblage of flora and fauna, combined with human interactions with the natural ecosystem, make this area socio-ecologically significant. In addition to being a protected area, Gosaikunda holds cultural and religious significance, with traditional indigenous peoples’ settlements within the park. Herding is a major livelihood practice among the local community, and transhumance is adopted for generations as a strategy for harnessing the resources among the herders (Aryal et al. 2014). The Gosikunda sector of LNP receives much higher rainfall (1972-2009 average annual rainfall at Dhunche, Rasuwa– the nearest metrological station at an elevation 2065 masl: ∼1885 mm) compared to drier rain-shadowed inner Himalayan Langtang-Kyangjing valley (average annual rainfall: ∼650 mm) (Chaudhary 1998; Tripathee et al. 2014).

**Fig. 1:**
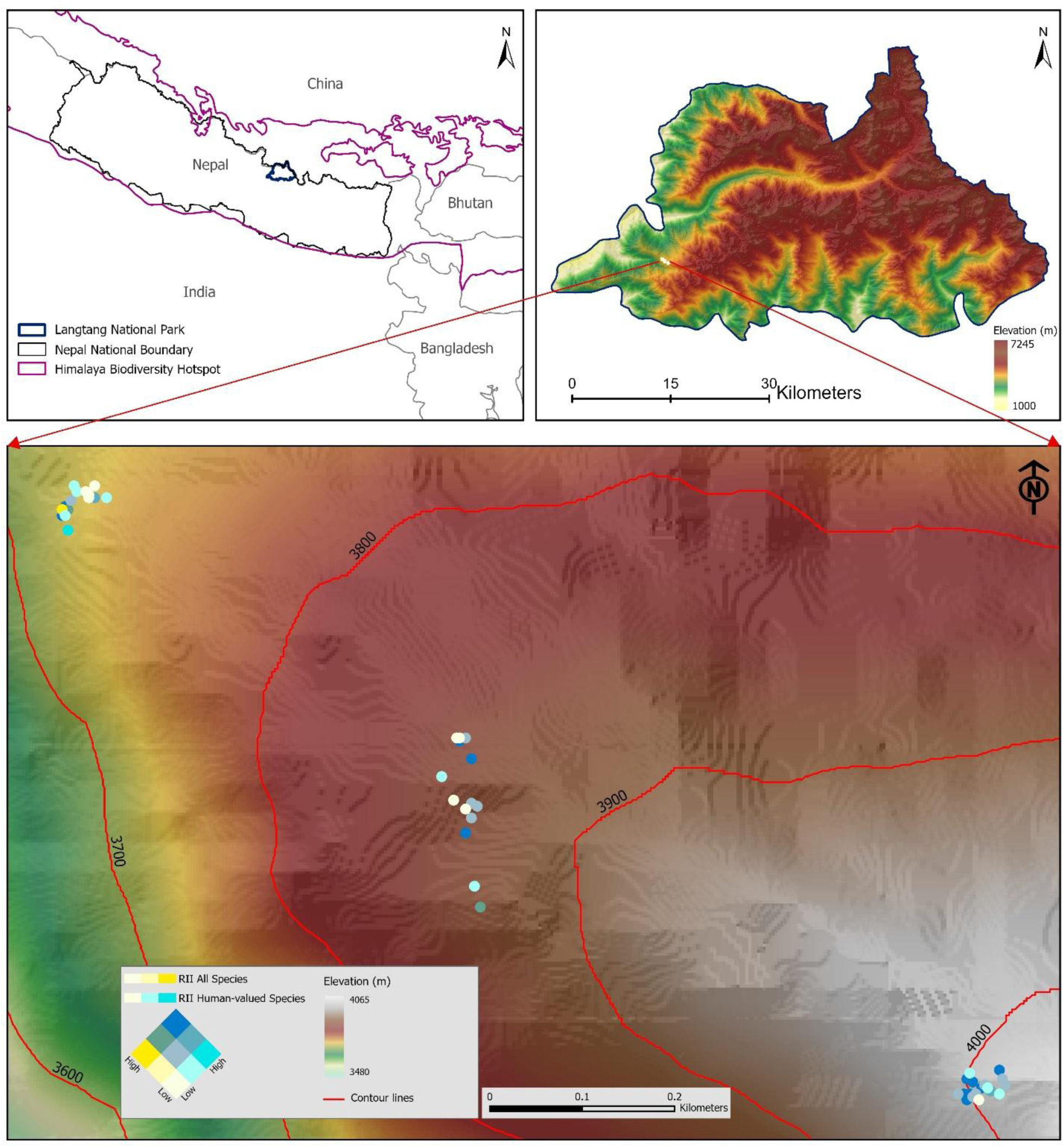
Study area map showing 21 paired plots sampled in each of three elevation belts. The bivariate colors symbolize the values of relative interaction intensity (RII) of both all and human-valued species, where dark blue represents plots with high-high values and light cream color represents low-low values. Note the greater number of dark blue plots around 4000 masl (high elevation), illustrating the stronger facilitation by *Berberis* shrubs for both overall and human-valued species at higher elevation.

### Focal species (nurse plant)

*Berberis angulosa* is a spiny dwarf shrub endemic to the Himalayas and is among the dominant patch-forming shrubs in the region with around 15% cover (Adhikari et al. 2012; Miehe et al. 2015; Parajuli et al. 2021). Distinguished due to its bright red and usually solitary fruit, *Berberis angulosa* is a low stature shrub up to two meters tall (Adhikari et al. 2012), even though the average height in this study was recorded to be 0.58±0.02 m. An erect and profusely branched from base, this shrub has coriaceous leaves with 1-3 spinulose teeth on each side of lamina, along with both central (1-3 cm) and lateral (0.5-1.5 cm) spines on the stem (Adhikari et al. 2012), thus largely preventing livestock browsing within its canopies (Parajuli et al. 2021); RP and KA personal observations). *Berberis angulosa* was selected because its mosaic of distinct patches interspersed with open gaps was well suited to the pairwise sampling design employed in this study. For a more in-depth botanical description of this shrub, we refer to Adhikari et al. (2012).

### Sampling design

Pairwise sampling was conducted in patches of *Berberis* shrubs and corresponding open gaps randomly selected in three altitudinal belts: low (∼ 3730 m), mid (∼ 3870 m), and high (∼4010 masl), from just above Cholangpati to Lauribina areas of the Gosainkunda sector in Langtang National Park, Central Nepal. In each elevational transect, we selected patches of *Berberis* shrubs with an adjacent open area without shrubs. A total of 21 pairs of 1×1 m^2^ shrubs and open plots were sampled at each of the three elevational belts, for a total of 63 paired plots (i.e., 21 plots × 2 habitats × 3 elevations = 126 plots altogether). The shrub plots were laid completely within the canopy of *Berberis* at least 0.5 meters from the edge, whereas the open plots were sampled at around one meter from the shrub edge in the open gaps to eliminate possible bias due to edge effects (Parajuli 2012; Parajuli et al. 2021). Each plot was further divided into four equally sized subplots to facilitate the estimation of % covers of plant functional groups within the sample plot and to assess grazing (signs of livestock browsing, hoofmarks and droppings) intensity (Parajuli 2012; Parajuli et al. 2021). The purpose of the sampling was to examine the spatial effect of the presence of *Berberis* shrubs along altitude-mediated environmental gradients. Geographic positioning system coordinates (GPS locations) were marked, one for each paired plot, to visualize the geospatial pattern of the shrub’s facilitative effect on understory species, including along the elevation gradients (Fig. 1).

All vascular species present within sampled plots were recorded. Species were identified onsite by consulting the field manuals (Polunin and Stainton 1984). Unidentified species were collected, and herbarium voucher specimens were prepared. All species were later identified and confirmed by expert plant systematists at the National Herbarium and Plant Laboratories (KATH). Nomenclature followed national checklists such as (Rajbhandari et al. 2017, 2019, 2021, 2022; Shrestha et al. 2022a; Rajbhandari et al. 2025). Geographic distributions in these publications, along with the plants of the world online database by Royal Botanic Gardens Kew (https://powo.science.kew.org/), were used to determine the endemic status of each species.

### Field data recording

In each of the sampling plots, all vascular plants were documented based on their presence or absence. The percent covers of plant functional groups such as herbs and grasses were also recorded separately based on visual estimates and logged on a scale of 0-100%. To maintain consistency, the same researcher estimated the cover % for all functional groups and throughout the study. Additionally, we also documented environmental variables like altitude, slope, and aspect for each plot. Litter thickness was measured in centimeters (later converted to meters) using a steel ruler. Similarly, the height of shrubs was recorded using measuring tape. Observations were made regarding browsing, trampling, signs of plant harvest, and traces of fires within each plot. Because harvesting and fires were infrequent, we removed those from analysis. A composite ‘grazing’ disturbance variable was determined based on signs of browsing and trampling on a scale of 0-4, where 0 is absence of grazing and 4 means heavy grazing (Parajuli et al. 2021).

Plants known to have medicinal values or other socio-cultural significance were considered as useful plants of humans (hereinafter human-valued species) following the same categories as in (Parajuli et al. 2021). Broadly, the species’ usefulness categories were medicinal, food, additive, social, commercial, and fodder values (see details in (Parajuli et al. 2021). The identification of plant species’ usefulness was determined in consultation with local peoples, herders and traditional healers (n=23) and by consulting relevant published sources (e.g., (Manandhar 2002; Baral and Kurmi 2006; Ghimire et al. 2008a). Verbal consent was obtained prior to the consultation and all local resource persons voluntarily shared their traditional knowledge about plant use values. The conservation status of species were based on (Ghimire et al. 2008a; Bachman et al. 2024).

### Statistical analysis

We employed the relative interaction indices as proposed by (Armas et al. 2004), to assess the extent of interaction between the shrub and different species. The index is calculated using the equation RII = (S_s_ – S_o_) / (S_s_ + S_o_), where S_s_ represents the number of species occurring within the shrub plot and S_o_ represents the number of species in the open plot. We calculated relative interaction indices for all species and human-valued species, i.e., RII of total species (RII_tsp_) and RII using a dataset with human-valued species only (RII_hvsp_). These indices were computed for each pair of sample plots.

To test the patterns of RIIs along elevation gradients we first fitted a linear regression model (LM) with the Gaussian family, because RIIs followed the normal distribution, using *lm()* function in the R statistical environment (R Core Team 2022). We also included other predictors such as litter thickness measured in shrub plots and grazing disturbance recorded in open areas. We tested heterogeneity using Levene’s test and fitted a generalized least square regression (GLS) model as an additional attempt to check for the potential heterogeneity inherently associated with ecological data using different variance structures (Zuur et al. 2009). Because both Levene’s test and the model comparisons using *anova()* were non-significant we ruled out the GLS. However, given the nested sampling design (21 pairs of plots sampled within each elevation belt), we also did fit linear mixed-effects models (LMMs) to analyze the variation of RII_tsp_ and RII_hvsp_ along the elevation gradients using *lme()* function from the *nlme* package (Pinheiro et al. 2023). Altitude (a factor with three levels: low, mid, and high), litter thickness and grazing (a factor with five levels: 0-4) were treated as a fixed factors, while plot (random intercept with 63 levels) was considered as a random term to account for variability in RII values for both total species and human-valued species across plots, which is specifically important to consider in the hierarchical sampling structure like this. The LMMs were fitted with ‘ML’ (maximum likelihood) and compared with fixed-effects only model, a common practice of model selection process suggested and applied in ecology (Zuur et al. 2009; Bolker et al. 2009).

The non-significant likelihood ratio test, along with higher AIC and BIC, indicated that LMMs didn’t perform better than LMs. Since there were no apparent signs of heterogeneity and both GLS and LMMs failed to improve the model fit over LMs, we decided to use linear models although all plots were not completely independent due to our nested sampling design. Because the overall effects of elevation, our fixed factor of primary interest, on RIIs were consistent regardless of the model used and LMs diagnostics showed the good sign of model fit (i.e., the residuals were normally distributed and fit vs residuals were random without any visible patterns), the conclusions from the LMs should be robust enough to draw overall patterns of RIIs along elevation gradients. Finally, we also compared the mean differences in RIIs computed with covers of herbs and grasses between the elevation bands using ANOVA and calculating the mean and 95% confidence intervals (CIs). All visual comparisons were made using point and whisker plots with mean and 95% CIs using the *ggplot2* package in the R statistical environment.

We conducted Nonmetric Multidimensional Scaling (NMDS) ordination using a Bray-Curtis dissimilarity matrix to visualize differences in species composition between areas under *Berberis* shrubs and adjacent open areas along an altitudinal gradient. We employed Permutational Multivariate Analysis of Variance (PERMANOVA) with 999 permutations to test for overall difference in species composition between habitat types (under shrubs vs. open) and altitudinal levels (low, mid, high). Similarly, we performed pairwise PERMANOVA to compare the compositional difference between each of the habitat types and elevation adopting a similar approach. To analyze the effect of grazing and litter content in species composition, we fitted these variables to NMDS objects using the *env_fit()* function in the *vegan* package (Oksanen et al. 2025) in R, with 999 permutations and created the plots with *ggplot2* package in R (R Core Team 2022).

### Indicator species analysis

We used indicator species analysis (ISA) to test the association of species with a habitat (shrubs vs open) and statistical significance was tested using a Monte Carlo 9999 permutation procedure using *multipatt()* function in R package *indicspecies* (De Cáceres and Legendre 2009). We used group equalized association function (Indicator Value Index, IndVal.g) method to identify whether a species has an ecological association with either shrubs or open areas. The association statistics in IndVal is a product of two quantities, namely ‘A’ and ‘B’ (Dufrêne and Legendre 1997; De Cáceres and Legendre 2009), where, for presence–absence data, component A is “the relative frequency of the species in the target site group divided by the sum of relative frequencies over all groups”, whereas B is “the relative frequency of occurrence of the species inside the target site group” (De Cáceres and Legendre 2009). The indicator species analysis is commonly used in facilitation and other community ecological research to identify most characteristic species representing habitat of interests using both presence-absence and abundance data (Dufrêne and Legendre 1997; De Cáceres and Legendre 2009; Parajuli et al. 2021).

### Phylogenetic diversity

We created a phylogenetic tree (Fig. 2) by grafting our list of the taxa into GBOTB.extended mega-tree using *V.PhyloMaker* package (Jin and Qian 2019) in R statistical environment. Mean Phylogenetic Distance (MPD), which is an average evolutionary distance between all pairs of species, is calculated as a measure of phylogenetic diversity. We estimated Net Relatedness Index (NRI) by comparing the observed value of MPD with a null model generated with 999 randomizations of the community matrix. We calculated NRI using the equation:

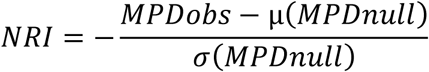

**Fig. 2:**
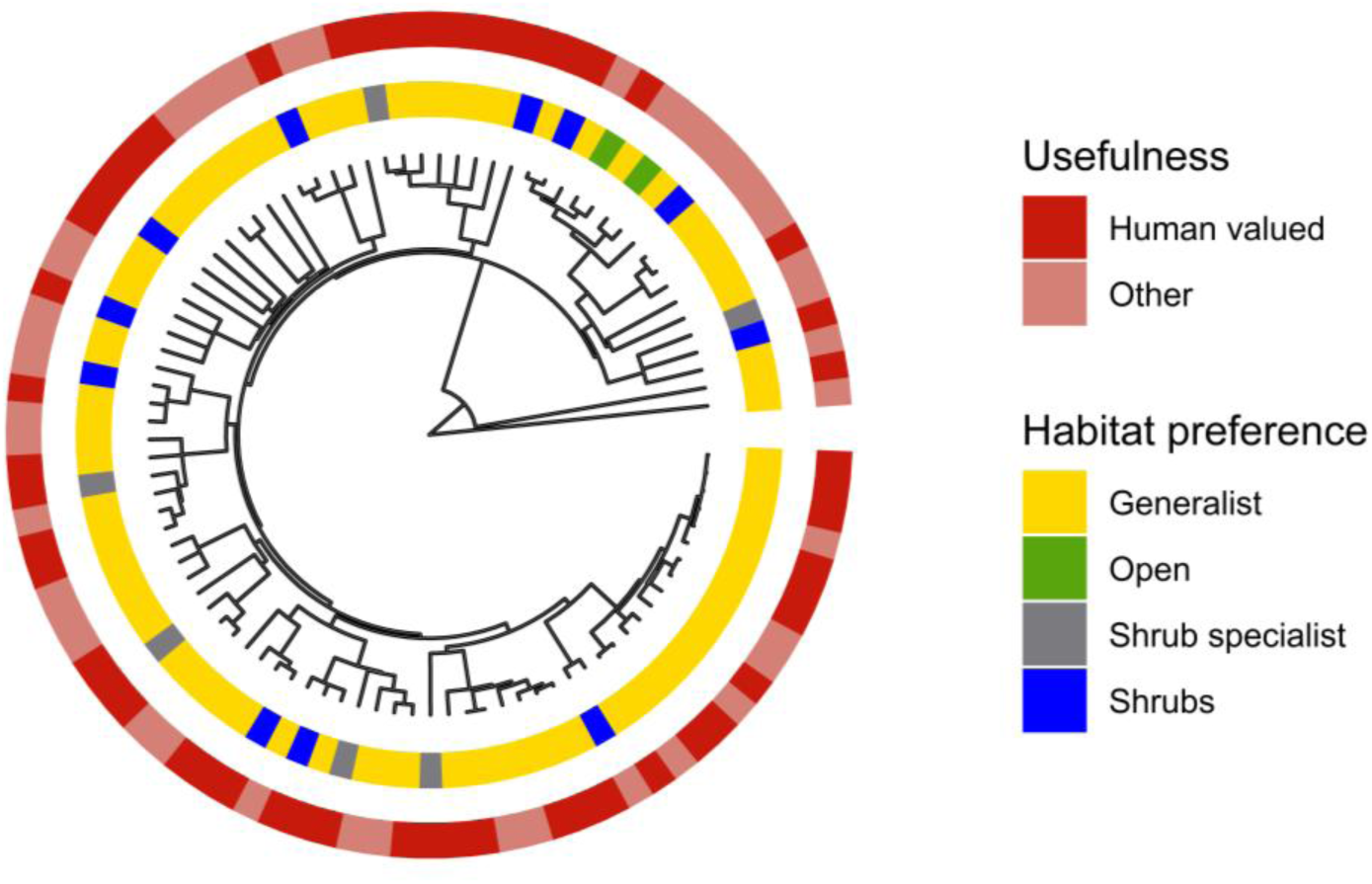
Phylogenetic tree of the taxa recorded in this study; highlighting the diversity in ecological, phylogenetic and the usefulness. The inner colored bar outside the tips level represents the habitat preference of species, and the outer circle indicates if the species has human-value highlighting phylogenetic and usefulness diversity. Where, Shrubs = species with significant association with shrubs but also present in open areas, Open = species with significant association with open areas but also present in shrubs plots, and Shrub specialists = species unique to *Berberis* patches and the association was statistically significant. Note that majority of species having significant association with *Berberis* are human-valued.

Where, MPDobs is the observed mean pairwise distance, μ(MPDnull) and σ(MPDnull) are the mean and standard deviation of MPD from randomized communities. The diversity matrices were calculated using *Picante* package (Kembel et al. 2020) in R statistical environment. ANOVA was done to find the difference between the NRI values along the elevation gradients and the habitat type. We conducted a two-way ANOVA to analyze the variation in NRI value in combination with habitat type and elevation. Post hoc analysis was done to check how NRI values differ in combination with each of the habitats and elevation belts.

## RESULTS

### Higher community and plot level richness beneath *Berberis*

A total of 97 vascular plant species were recorded in this study (Supplementary Information, SI, Table S3). Of these, 39 species were exclusively recorded from underneath the *Berberis* canopies, whereas only two species were confined to open areas. Similarly, 53 (out of 97) species were documented to be human valued species, which included 19 species unique to shrubs plots compared to only one from open gaps. The mean richness of both total species and human-valued species were significantly higher within shrub canopies than in adjacent open gaps (total species: *t* = 14.56, *p* < 0.001; human-valued species: *t* = 9.5, *p* < 0.001). On average, shrubs plots have greater than 50% more species (mean richness of total species: 16.05, 95% CIs 15.2-16.9; human-valued species: 11.03, 95% CIs 10.36-11.7) than in open plots (mean richness of total species: 10.25, 95% CIs 9.7-10.8; human-valued species: 6.9, 95% CIs 6.5-7.3). Likewise, the % cover of herbaceous plants was significantly higher underneath *Berberis* (mean = 58.94, 95% CIs 53.4-64.5) than in open (mean = 47.48, 95% CIs 43.2-51.7), whereas grass coverage was higher in open (mean = 8.68, 95% CIs 6.9-10.4) than in shrub plots (mean = 3.58, 95% CIs 2.5-4.8).

### Increase in relative interactions intensity along elevation gradients

The average RII of both all and human-valued species were positive and significantly different from zero (RII_tsp_: mean = 0.22, 95% CIs = 0.19–0.25; RII_hvsp_: mean = 0.22, 95% CIs = 0.19–0.26) suggesting a facilitative role of *Berberis* shrubs on understory species in Gosaikunda sector of LNP, Nepal. Our test of trends of facilitatory effects of *Berberis* shrubs along the elevation gradients showed an interesting pattern, albeit the facilitation intensity was the strongest at the highest elevation for both all and human-valued species. The difference in RII_tsp_ was statistically significant between low vs high elevation, however, it was non-significant when compared mid-elevation with both low and high elevations (Fig. 3; SI Table S1). The pattern was similar for human-valued species (RII_hvsp_) too.

**Fig. 3:**
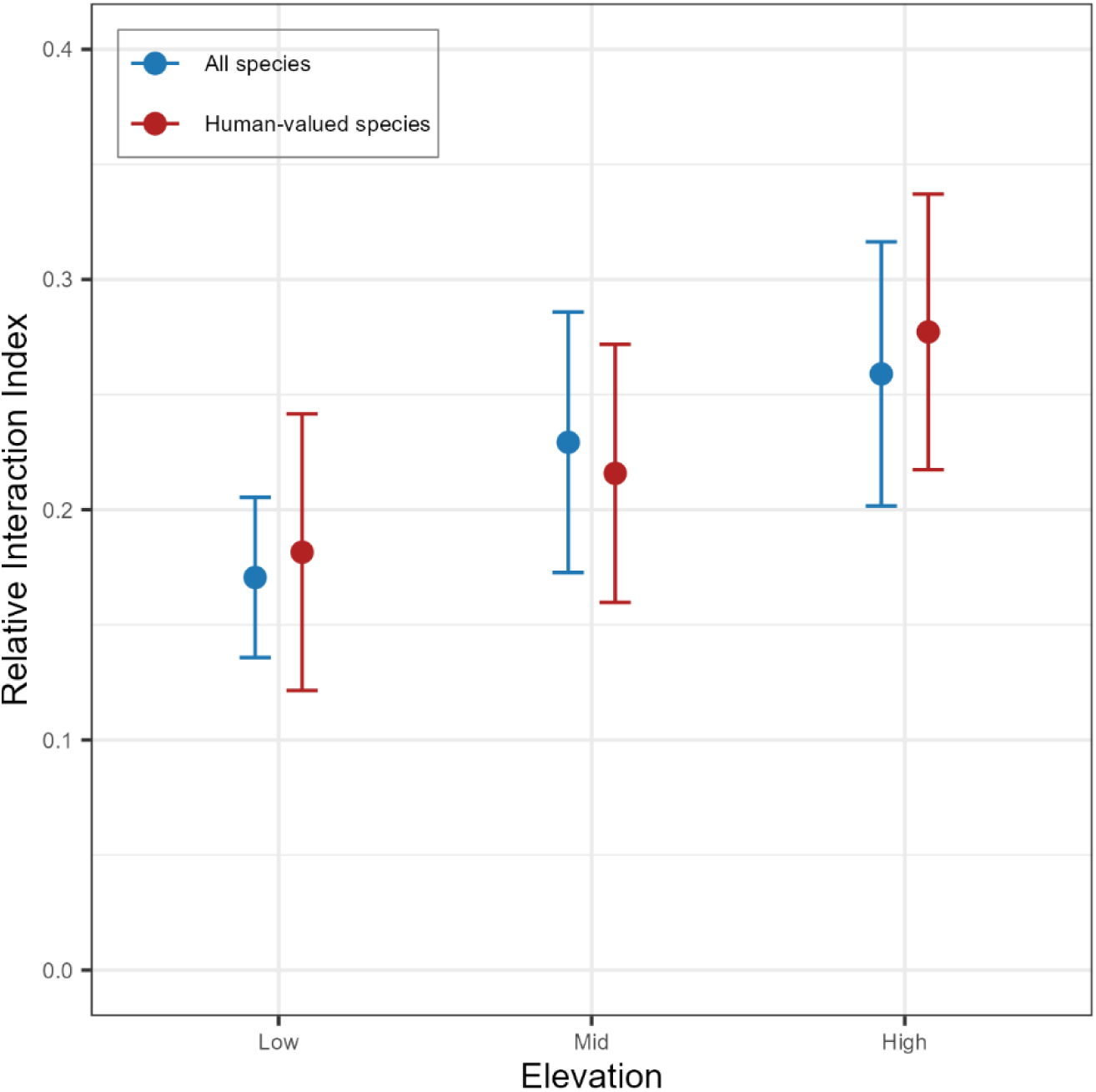
Dot and whisker (mean±95% CIs) plots showing the patterns of interactions between *Berberis* shrubs and understory species, both all (RII_tsp_) and human-valued (RII_hvsp_), along elevation gradients. The highest intensity of facilitations by *Berberis* shrubs for both total and human-valued species were evident at high-elevation.

The mean RIIs values of % covers of herbs and grasses revealed varied patterns in their interactions with *Berberis* shrubs. Overall, as illustrated by the RII computed with % covers, herbaceous plants (mean = 0.09, 95% CIs = 0.02–0.16) had significantly positive interactions with *Berberis* shrubs, whereas the grasses exhibited strong competition (mean = -0.36, 95% Cis = -0.48 – -0.24). Per elevation-wise comparisons, while the facilitative effects of shrubs on covers of herbs and the competitive effects on grass were strongest at high-elevation, these interactions were neutral, i.e. non-significantly different from zero, at low-elevation (Fig. 4).

**Fig. 4:**
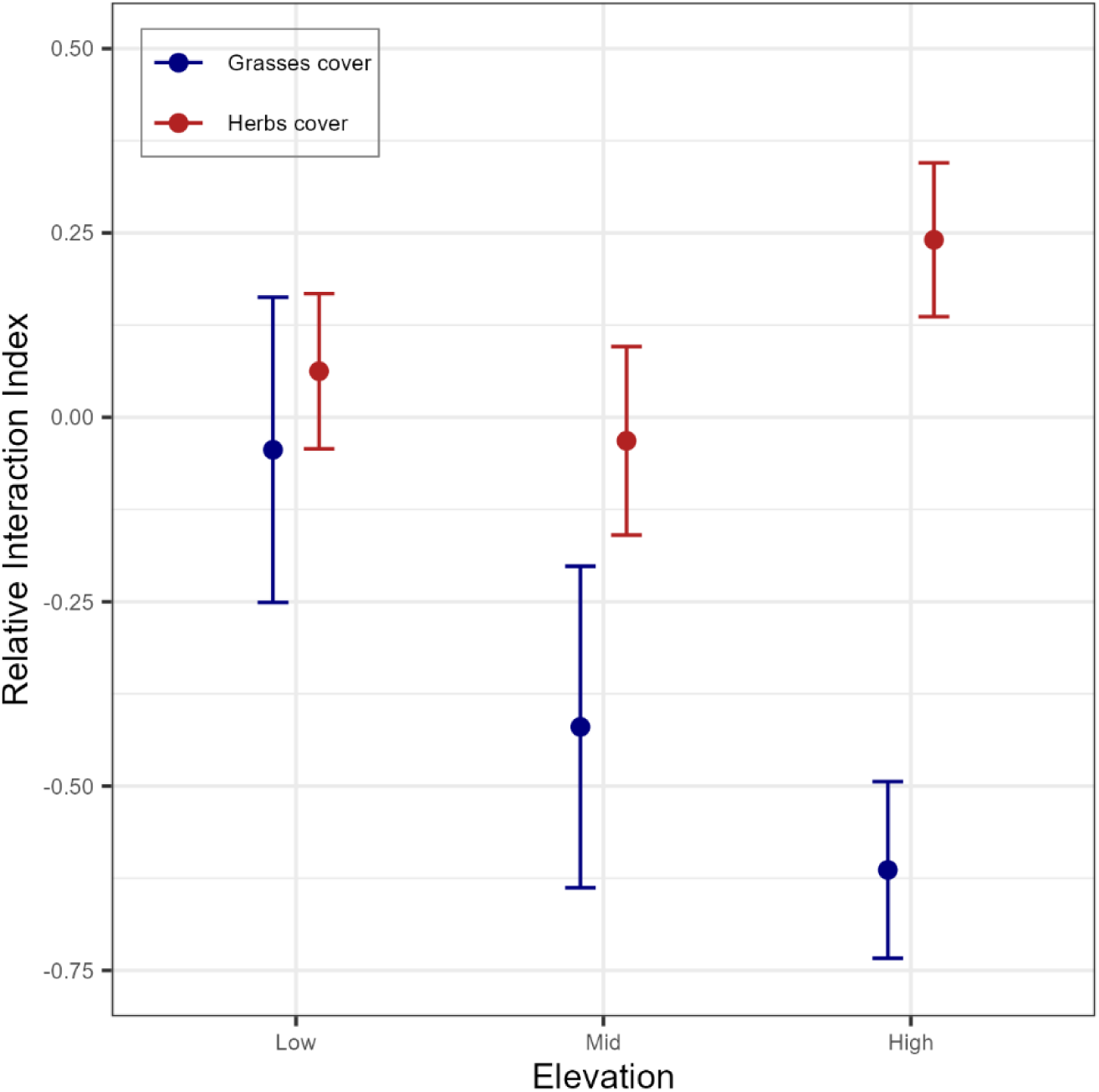
Overall trends of RIIs computed with % covers of different functional groups along elevation gradients. Note the significantly strong positive interactions between *Berberis* shrubs with herbs at high-elevation, non-significant in the case of grasses and herbs at low-elevations, and significantly negative interaction (i.e., competition) with grasses at mid- and high-elevations.

### Difference in species composition between shrubs and open habitats

Overall, species composition differed significantly between plots in open areas and under shrubs for both all species (*F* = 24.77, *p* = 0.001) and human valued species (*F* = 29.21, *p* = 0.001). It varied significantly between open gaps and shrub canopies across all three elevations as well for both all (*F* = 21.96, *p* = 0.001; Fig. 5A) and human-valued species (*F* = 26.41, *p* < 0.001; Fig. 5B). Post-hoc pairwise comparison between the habitat types at low, mid, and high elevations also showed significant difference in composition for both all species (*F* = 17.23, *p_adj_* = 0.01; *F* = 16.32, *p_adj_* = 0.01; *F* = 21.0, *p_adj_* = 0.01) and human-valued species (*F* = 22.79, *p_adj_* = 0.01; *F* = 17.76, *p_adj_* = 0.01; *F* = 25.31, *p_adj_* = 0.01). Community composition in open plots was significantly correlated with grazing disturbance (all species: *R²* = 0.42, *p* = 0.001; human-valued: *R²* = 0.42, *p* = 0.001), while in shrub plots it was significantly correlated with litter thickness (all species: *R²* = 0.57, *p* = 0.001; human-valued: *R²* = 0.55, *p* = 0.001).

**Fig. 5:**
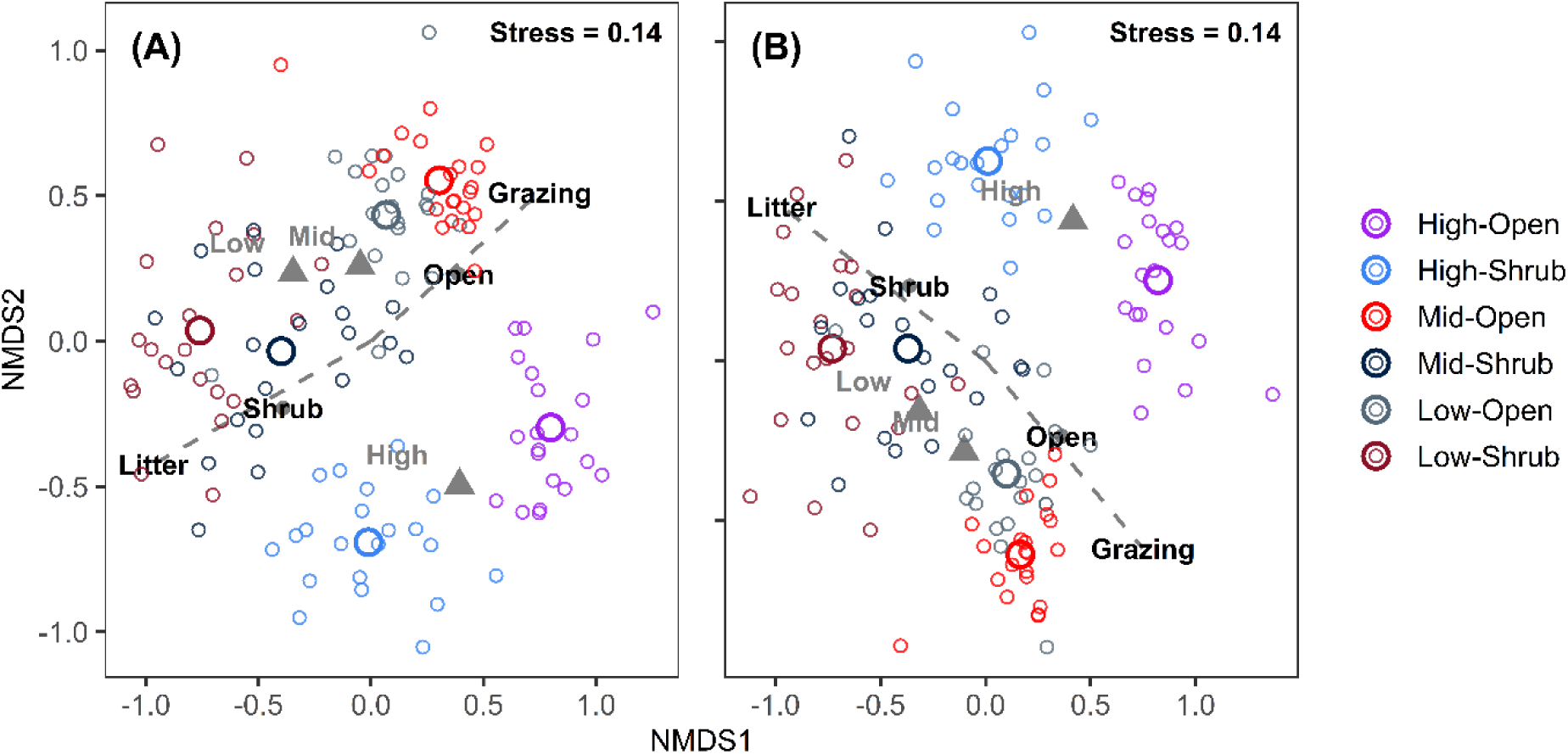
NMDS plots illustrating differences in taxonomic composition between open and shrub habitats across the altitudinal range for (A) all species and (B) human-valued species. Small colored circles represent plot-level values and bigger colored circles represent the centroid of habitat types in different elevations. Grey triangles are the centroids of pooled (shrubs + open) species composition at three different elevations.

### More high value plants associated with shrubs patches than open gaps

We found around one third of characteristic species (30 out of 97) that had statistically significant association with either of the habitats (Table 1). A total of 25 species (i.e., 83%) had significant association with shrubs compared to only five (i.e., 17%) species in the case of open areas. Of these, 12 species were shrub specialists, i.e., species unique to *Berberis* shrubs with their association being statistically significant, whereas none of the species were found to be open specialists. Likewise, the proportion of human-valued species having significant association with shrubs was much higher (15 out of 17) than those associated with open areas (2 out of 17). Some commercially important medicinal herbs, e.g., *Delphinium himalayae* (Nationally and globally threatened, Himalayan endemic), *Rheum acuminatum* (Himalayan endemic), and *Valeriana hardwickii*, were shrub specialists and hence protected within the patches of *Berberis*. Some other high-value medicinal plants with conservation importance such as, *Allium wallichii* (Nationally threatened), *Parnassia nubicola* (Locally vulnerable), *Aconitum ferox* (Nationally and globally threatened, Himalayan endemic), though were less frequent and statistically none-significant, were only recorded from underneath *Berberis* shrubs.

**Table 1:** Results of test of species’ habitat association with shrubs and open areas. Only species with statistically significant association with either of the habitats were presented here. Association statistics are the indicator value test statistics obtained from ISA using *IndVal.g* function and *p*-values were generated by Monte Carlo permutation procedure with 9999 random resampling. Species used in various local indigenous traditional practices are marked as ‘Human valued’. Habitat association terms, Shrubs = species with significant association with shrubs but also present in open areas, Open = species with significant association with open areas but also present in shrubs plots, and Shrub specialists = species unique to *Berberis* patches and the association was statistically significant. A full list of species with their association statistics and corresponding *p*-values is given in SI Table S2.

| *Species name* | Association Statistics | *P*-value | Habitat Association | Usefulness | Endemism | Conservation Status |
| --- | --- | --- | --- | --- | --- | --- |
| *Saxifraga diversifolia* | 0.77 | 0.0001 | Shrubs | Human valued |  |  |
| *Potentilla daltoniana* | 0.77 | 0.0001 | Shrubs | Human valued |  |  |
| *Themeda triandra* | 0.66 | 0.0001 | Shrubs | Human valued |  |  |
| *Calamagrostis pseudophragmites* | 0.65 | 0.0001 | Open |  |  |  |
| *Anaphalis nepalensis* | 0.65 | 0.0001 | Shrubs | Human valued |  |  |
| *Corydalis juncea* | 0.59 | 0.0001 | Shrubs | Human valued | Himalayan endemic |  |
| *Stellaria patens* | 0.58 | 0.0001 | Shrubs |  | Himalayan endemic |  |
| *Valeriana hardwickii* | 0.53 | 0.0001 | Shrub specialist | Human valued |  |  |
| *Impatiens* sp. | 0.47 | 0.0001 | Shrubs |  |  |  |
| *Tetrataenium wallichii* | 0.47 | 0.0001 | Shrub specialist | Human valued | Himalayan endemic | Threatened |
| *Cyperus* sp. | 0.49 | 0.0002 | Open |  |  |  |
| *Cyanaea* sp. | 0.44 | 0.0002 | Shrub specialist |  |  |  |
| *Caltha palustris* | 0.53 | 0.0004 | Open | Human valued |  |  |
| *Gentiana depressa* | 0.51 | 0.0004 | Shrubs | Human valued |  |  |
| *Hemiphragma heterophyllum* | 0.44 | 0.0004 | Shrub specialist | Human valued |  |  |
| *Pedicularis megalantha* | 0.48 | 0.0006 | Shrubs | Human valued | Himalayan endemic |  |
| *Cyananthus himalaicus* | 0.47 | 0.0006 | Shrubs |  | Nepal endemic | Threatened |
| *Euphorbia stracheyi* | 0.46 | 0.0011 | Shrubs | Human valued |  |  |
| *Aster* sp. | 0.40 | 0.0014 | Shrub specialist |  |  |  |
| *Androsace sarmentosa* | 0.40 | 0.0024 | Shrub specialist | Human valued | Himalayan endemic |  |
| *Sporobolus fertilis* | 0.49 | 0.0029 | Open | Human valued |  |  |
| *Epilobium* sp. | 0.38 | 0.0029 | Shrub specialist |  |  |  |
| *Galium* sp. | 0.38 | 0.0034 | Shrub specialist |  |  |  |
| *Delphinium himalayae* | 0.36 | 0.0055 | Shrub specialist | Human valued | Himalayan endemic | Threatened |
| *Calamagrostis scabrescens* | 0.41 | 0.0130 | Shrubs |  |  |  |
| Unknown1 | 0.33 | 0.0131 | Shrub specialist |  |  |  |
| *Arisaema jacquemontii* | 0.38 | 0.0136 | Shrubs | Human valued |  |  |
| *Rheum acuminatum* | 0.33 | 0.0138 | Shrub specialist | Human valued | Himalayan endemic |  |
| *Plantago* sp. | 0.38 | 0.0178 | Open |  |  |  |
| *Arisaema* sp1 | 0.31 | 0.0270 | Shrub specialist |  |  |  |

### Higher phylogenetic diversity under the shrub canopy

Elevation significantly affected NRI for all species (*F* = 5.65, *df* = 2, *p* < 0.001) and human-valued species (*F* = 13.08, *df* = 2, *p* < 0.001) (Fig. 6). Post-hoc comparison showed that, for all species, NRI differed significantly only between low and high elevations (*p* < 0.003), whereas in the case of human-valued species, significant differences were observed between low and high (*p* < 0.001) and mid and high elevations (*p* < 0.001). Similarly, habitats have had significant effects on NRI for all species (*F* = 20.18, *df* = 1, *p* < 0.001) and human-valued species (*F* = 16.82, *df* = 1, *p* < 0.001). Compared to open gaps shrubs had significantly lower NRI for both overall (shrubs: mean = -0.49, 95% CIs -0.71 – -0.27; open: mean = 0.12, 95% CIs -0.04 – 0.28) and human-valued species (shrubs: mean = -0.65, 95% CIs -0.82 – -0.49; open: mean = 0.49, 95% CIs 0.25 – 0.73). The interaction between habitat and elevation was also significant for all species (*F* = 3.94, *df* = 2, *p* = 0.02) and human-valued species (*F* = 3.18, *df* = 2, *p* = 0.04) (Fig. 6). However, post-hoc analysis revealed that the difference in NRI values between shrubs vs open habitats was significant only at mid elevation for all species (*p_adj_* < 0.001), whereas, at all three elevations in the case of human values species (*p_adj_* < 0.001 for all three elevations) (Fig. 6).

**Fig. 6:**
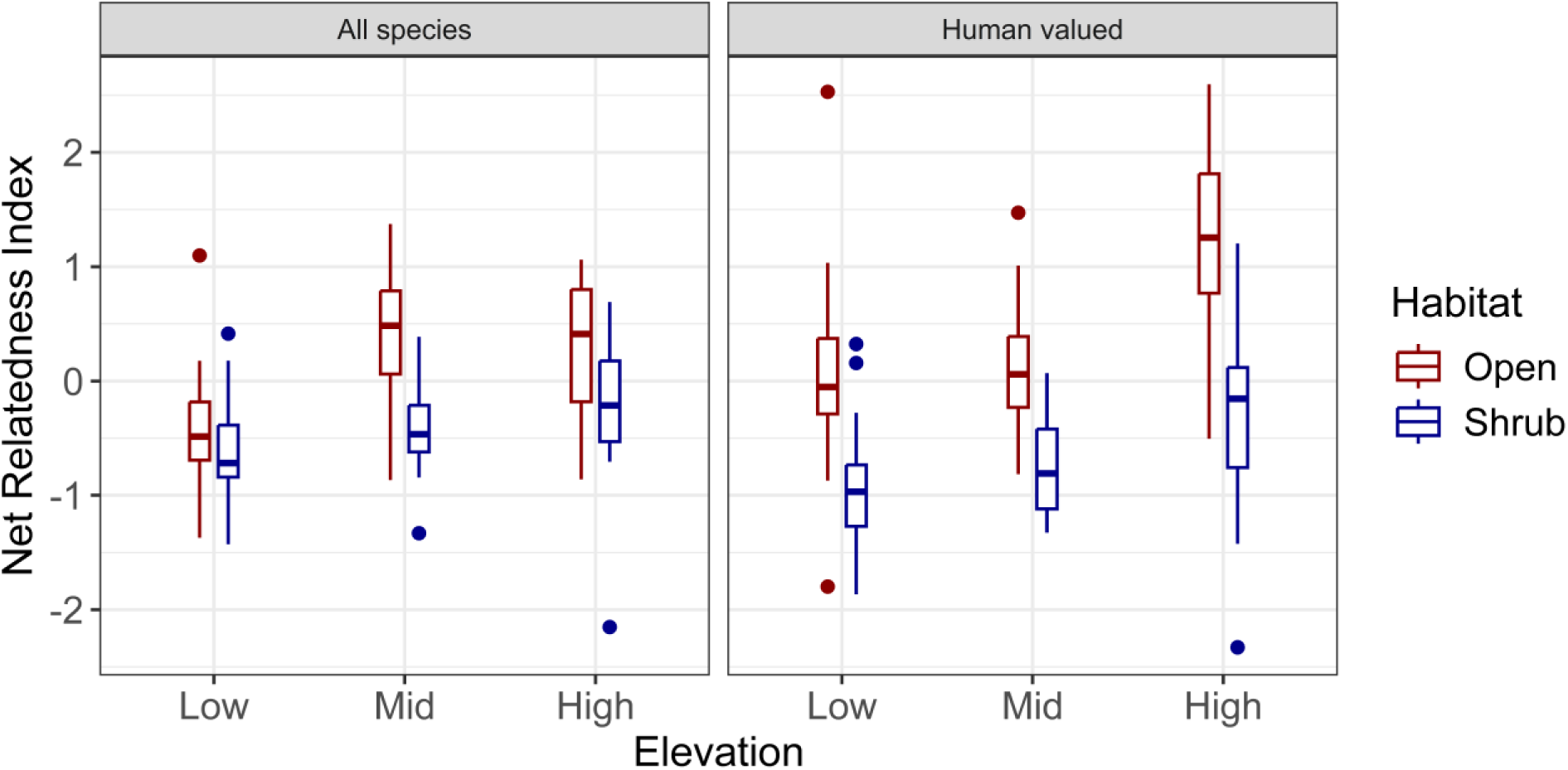
Comparison of NRI values between open and shrub habitats across the altitudinal range for all species and human-valued species, showing the lower median NRI values in shrubs at all three elevations.

## DISCUSSION

Our results demonstrate that facilitation by *Berberis* plays a critical role in promoting the taxonomic and phylogenetic diversity of alpine plants in Nepal’s Himalaya. Previous studies also reported a strong role of facilitation by shrubs and cushion plants in maintaining plant diversity in the Himalayas (e.g., (Yang et al. 2010; Chen et al. 2015; Ale et al. 2018, 2023; Parajuli et al. 2021); but see (Dvorský et al. 2013; Ram and Chawla 2024). Although the current research site is a relatively wetter Himalayan region compared to the dry inner Kyangjing valley of Langtang National Park in (Parajuli et al. 2021), collectively these studies provide consistent evidence of the facilitative role that nurse shrubs play in structuring plant communities in the high-elevation mountains regardless of difference in precipitation regimes. Furthermore, our findings support the idea of nurse plant-created microsites as conservation refugia of plant taxonomic and phylogenetic biodiversity in the Himalayas (Parajuli et al. 2021; Rasray et al. 2024). Moreover, this study may be the first to demonstrate that facilitation by shrubs contributes to preserving phylogenetic diversity of high-value medicinal and other species of societal and conservation importance.

We observed the strongest facilitation by *Berberis* at the highest elevation for both the overall plant community and for human-valued species. These findings align with the predictions of the Stress Gradient Hypothesis (SGH; (Bertness and Callaway 1994) and are consistent with reports from previous studies conducted in the Himalayan region (e.g., (Yang et al. 2010; Ale et al. 2018; Parajuli et al. 2021); but see (de Bello et al. 2011; Dvorský et al. 2013). In the Himalayas and other alpine mountain systems, environmental stress typically increases with an increase in elevation due to declining temperature, shortening growing seasons and increasing exposure to radiation (Körner 2007, 2021; Sundqvist et al. 2013). For instance, the average temperature lapse rate in Nepal’s Himalaya is -0.5°C to -0.7°C per 100 meters of elevation gain (Kattel et al. 2013; Khadka et al. 2023). Given the elevation difference of approximately 300 m between the low- and high-elevation transects in our study, this translates to decline of about 2°C in mean surface air temperature, resulting in more stressful climatic conditions at higher elevations. These increments in climatic and other environmental severities could explain the strongest facilitation observed at the high-elevation sites, in line with SGH predictions (Bertness and Callaway 1994). Notably, with only a 200 m elevation gain, (Yang et al. 2010) documented a stronger facilitative effect at higher site (4700 m) compared to the lower one (4500 m) in the Hengdung alpine mountains of the Sino-Himalayas, underscoring how small changes in elevation can influence plant-plant interactions patterns in these youngest mountain systems.

Per our expectations, unique species were observed in both habitats, albeit the proportion was much higher underneath *Berberis* canopies (40.2% of total species, 35.9% of total human-valued) than open gaps (2.1% of total species, 1.9% of total human-valued), leading to a significant difference in community compositions of both overall and human-valued species between these two contrasting habitats. Since livestock browsing, as a part of centuries long transhumance agropastoral practice, is common in Nepal’s Himalayan landscapes including Langtang region (Aryal et al. 2014, 2015), largely palatable yet grazing intolerant species were benefitted from protection by the patches of *Berberis* (Parajuli et al. 2021); RP and KA personal observations and interactions with local herders), an erect thorny dwarf shrub (Adhikari et al. 2012). Nurse shrubs, specifically spiny ones, are generally found to be effective in protecting palatable species in heavily grazed landscapes (Howard et al. 2012; Tirado et al. 2015; Parajuli et al. 2021). The indicator species of open areas were mostly grazing-tolerant but shade-intolerant perennial grasses, such as *Calamagrostis pseudophragmites, Sporobolus fertilis, Cyperus* sp., *Kobresia* sp. or generally undesirable to livestock or unpalatable forbs, e.g., *Caltha palustris, Plantago* sp. (Aryal et al. 2015; Paudel et al. 2020; Parajuli et al. 2021); RP and KA personal observations and interactions with local herders). This could be the reason for a strong competitive interaction between grasses and Berberis as shown by significantly negative RII (mean = -0.36, 95% CIs = -0.48 – -0.24) when computed with % cover values of grasses. In contrast, *Berberis*-specialist species were dominantly forbs with low to moderate tolerance to livestock browsing and highly valued because of their medicinal and other societal uses, endemism and conservation status such as *Delphinium himalayae, Rheum acuminatum Tetrataenium wallichii, Valeriana hardwickii* (Ghimire et al. 2008a; Paudel et al. 2020; Parajuli et al. 2021); RP and KA personal observations and interactions with local healers).

Our findings showed clear phylogenetic diversification within shrubs for both all species and the human-valued species recorded in the study area. *Berberis* supports communities with diverse evolutionary relationships, with human-valued taxa showing greater phylogenetic diversity within the shrub patches. In contrast, open areas were characterized by taxa with more similar evolutionary histories. This pattern was consistent across elevations. Nurse plants are known to promote phylogenetic diversity in environmentally severe ecosystems such as alpine, arid and dryland sites on several continents (Pistón et al. 2016; Pashirzad et al. 2019; Bashirzadeh et al. 2022), as well as degraded lands like mining fields in Spain (Navarro-Cano et al. 2016). *Berberis* is recognized for its botanical nursing effects enhancing species richness in stressful environments of the Himalayas, including more sensitive subsets of plant biodiversity such as the human-valued commercial and conservation priority species (Parajuli et al. 2021). Our finding evidenced that facilitation by nurse shrubs is equally effective in preserving phylogenetic diversity as well, implying its critical role in conserving evolutionary legacies and adaptations unique to the Himalayan environments.

Habitat patches created by *Berberis* tend to be comparatively nutrient rich with higher litter accumulation beneath their canopies (Parajuli et al. 2021), thus preserving soil moisture content and maintaining nutrient cycling (Callaway 2007). By creating such a microenvironment that can facilitate the growth and survival of unique assemblage of species, *Berberis* promotes phylogenetic diversity in high-elevation mountains that experience a variety of abiotic and biotic stressors (Körner 2021). Phylogenetic dispersion reflects the assembly of taxa with different traits over time (Webb et al. 2002). Taxa recorded from outside of *Berberis* patches were adapted to generally uniform habitat conditions common in open areas across landscape such as low nutrient availability and high grazing pressure (Parajuli et al. 2021); RP and KA personal observations), resulting them to be more similar in their traits due to habitat filtering (Webb et al. 2002, 2006). Whereas nurse shrubs facilitate species coexistence (Callaway 2007; Bulleri et al. 2016) and conserve regeneration niche of distantly related species that support higher phylogenetic diversity (Valiente-Banuet and Verdú 2007; Pistón et al. 2016).

Since at a global scale cushions plants found to serve as micro-refugia by facilitating less stress-tolerant lineages in severe alpine environments (Butterfield et al. 2013), the phylogenetically diverse assemblage of species, including high value plants, beneath *Berberis* confirms that nurse shrubs perform that important role in the Nepal’s Himalayas. Maintaining phylogenetic diversity is crucial for conserving evolutionary traits that are suited to diverse microclimates and, importantly, can withstand adverse conditions prevalent in the high mountain ecosystems (Faith 1992; Körner 2021). Moreover, preserving high phylogenetic diversity, specifically those of the habitat specialists, medicinal, endemic and endangered species that tend to have narrow niche space (Ahmad et al. 2021; Shrestha et al. 2022b), is critically important in the context of ongoing climate change and its some of the worsts impacts experienced in the Himalayas (Xu et al. 2009; Shrestha et al. 2012; Ahmad et al. 2025). Additionally, high-value medicinal and aromatic plants in Nepal’s Himalayas are experiencing extreme pressure due to overexploitation triggering population decline and increased extinction risk (Shrestha et al. 2022b; Khakurel et al. 2024).

Our findings clearly show that facilitation by nurse shrubs functions as a nature-based approach to conserve valuable plant biodiversity, while also opening new avenues to investigate how this ecological phenomenon helps sustain the human-ecosystem relationships that have been established in mountain environments for centuries. A phylogenetically diverse community not only preserves evolutionary history and adaptive potential, which are important for maintaining resilient ecosystems in rapidly changing environments (Faith 1992) but also maintains known and unknown biodiversity functions that may hold ecological or utilitarian values which can be of provisioning or cultural services to human beings (Cardillo 2023). *Berberis*, therefore, by protecting phylogenetically diverse taxa under its canopy, while conserving the evolutionary history shaped by plant-environment interactions in the Himalayan region is also maintaining the human-nature interactions in cultural landscapes such as those found in Langtang National Park.

## Acknowledgement

We would like to sincerely thank the Department of National Park and Wildlife Conservation (DNPWC), Kathmandu and LNP Headquarter, Rasuwa for granting the research permit to conduct field work and collect samples inside the national park. Similarly, we express our gratitude to local peoples, traditional healers and herders of Singompa, Thulo Bharkhu, Thulo Syabru, Cholangpati and Lauribina areas of LNP for sharing their knowledge on use of plants. We extend our sincere thanks to Robin Sarah Taylor for her support in grant proposal writing.

