## Supplementary Information for "Botanical nursing by a dwarf shrub as a nature-based approach to plant biodiversity conservation in the Nepal’s Himalayas"

Table S1: Linear model results summaries of RII for species

Table S2: Indicator Species Analysis full results

Table S3: Detail list of plants recorded and their use value

**Table S1: Linear model results summaries of RII for species**

**a) Summary table**

| a) All species | | | | b) Human-valued species | | |
|---|---|---|---|---|---|---|
| | Estimate | SE | t | Estimate | SE | t |
| Intercept | 0.252 | 0.049 | 5.05*** | 0.239 | 0.056 | 4.31*** |
| Elevation:low | -0.092 | 0.039 | -2.30 * | -0.116 | 0.045 | -2. 6* |
| Elevation:mid | -0.023 | 0.041 | -0.54 | -0.057 | 0.046 | -1.24 |
| Litter | -0.005 | 0.019 | -0.25 | -0.002 | 0.02 | -0.08 |
| Grazing1 | 0.018 | 0.049 | 0.36 | 0.059 | 0.056 | 1.06 |
| Grazing2 | 0.009 | 0.048 | 0.21 | 0.025 | 0.05 | 0.48 |
| Grazing3 | 0.015 | 0.054 | 0.28 | 0.064 | 0.06 | 1.06 |
| Grazing4 | 0.078 | 0.079 | 0.98 | 0.188 | 0.089 | 2.11* |

SE, Standard error of estimate, t, t-statistic; $^{\dagger}P<0.1$, *P < 0.05, **P < 0.01, ***P < 0.001

**b) ANOVA table**

| a) All species | | | b) Human-valued species | |
|---|---|---|---|---|
| | Df | F | Df | F |
| Elevation | 2 | 3.21* | 2 | $2.98^{\dagger}$ |
| Litter | 1 | 0.05 | 1 | 0.02 |
| Grazing | 4 | 0.26 | 4 | 1.31 |

D.f., degrees of freedom; F, F-statistic; $^{\dagger}P<0.1$, *P < 0.05, **P < 0.01, ***P < 0.001

**Table S2: Indicator Species Analysis full results**

A full list of tests of species' habitat association with shrubs and open areas along with association statistics and corresponding *p*-values. Association statistics are the indicator value test statistics obtained from ISA using IndVal.g function and *p*-values were generated by Monte Carlo permutation procedure with 9999 random resampling. Species used in various local indigenous traditional practices are marked as 'Human valued'. Habitat association terms, Shrubs = species with significant association with shrubs but also present in open areas, Open = species with significant association with open areas but also present in shrubs plots, and Shrub specialists = species unique to *Berberis* patches and the association was statistically significant, Shrubs only – NS = species present in shrubs plots only but the association statistically non-significant, Open only – NS = species recorded from open plots only but the association statistically non-significant, and Generalists = species without any habitat preferences, rather fairly commonly recorded from both habitats.

| Species name | Quantity 'A' | | Quantity 'B' | | | Association Statistics | *P*-value | Habitat Association | Usefulness | Endemism | Conservation Status |
|---|---|---|---|---|---|---|---|---|---|---|---|
| | Shrubs | Open | Shrubs | Open | Overall | | | | | | |
| Saxifraga diversifolia Wall. ex Ser. | 0.85 | 0.15 | 0.70 | 0.13 | 0.41 | 0.77 | 0.0001 | Shrubs | Human valued | | |
| Potentilla daltoniana (J. Gay) Mabb. | 0.81 | 0.19 | 0.73 | 0.17 | 0.45 | 0.77 | 0.0001 | Shrubs | Human valued | | |
| Themeda triandra Forssk. | 0.94 | 0.06 | 0.46 | 0.03 | 0.25 | 0.66 | 0.0001 | Shrubs | Human valued | | |
| Calamagrostis pseudophragmites (Haller f.) Koeler | 0.14 | 0.86 | 0.08 | 0.49 | 0.29 | 0.65 | 0.0001 | Open | | | |
| Anaphalis nepalensis (Spreng.) Hand.-Mazz. | 0.82 | 0.18 | 0.51 | 0.11 | 0.31 | 0.65 | 0.0001 | Shrubs | Human valued | | |
| Corydalis juncea Wall. | 0.84 | 0.16 | 0.41 | 0.08 | 0.25 | 0.59 | 0.0001 | Shrubs | Human valued | Himalayan endemic | |
| Stellaria patens D. Don | 0.92 | 0.08 | 0.37 | 0.03 | 0.20 | 0.58 | 0.0001 | Shrubs | | Himalayan endemic | |
| Valeriana hardwickii Wall. | 1 | 0 | 0.29 | 0 | 0.14 | 0.53 | 0.0001 | Shrub specialist | Human valued | | |
| Impatiens sp. | 0.94 | 0.06 | 0.24 | 0.02 | 0.13 | 0.47 | 0.0001 | Shrubs | | | |
| Tetrataenium wallichii (DC.) Manden. | 1 | 0 | 0.22 | 0 | 0.11 | 0.47 | 0.0001 | Shrub specialist | Human valued | Himalayan endemic | Threatened (Bachman et al., 2024) |
| Cyperus sp. | 0.06 | 0.94 | 0.02 | 0.25 | 0.13 | 0.49 | 0.0002 | Open | | | |
| Unknown2 | 1 | 0 | 0.19 | 0.00 | 0.10 | 0.44 | 0.0002 | Shrub specialist | | | |

| Species name | Quantity 'A' | | Quantity 'B' | | | Association Statistics | *P*-value | Habitat Association | Usefulness | Endemism | Conservation Status |
|---|---|---|---|---|---|---|---|---|---|---|---|
| | Shrubs | Open | Shrubs | Open | Overall | | | | | | |
| Caltha palustris L. | 0.13 | 0.87 | 0.05 | 0.32 | 0.18 | 0.53 | 0.0004 | Open | Human valued | | |
| Gentiana depressa D. Don | 0.83 | 0.17 | 0.32 | 0.06 | 0.19 | 0.51 | 0.0004 | Shrubs | Human valued | | |
| Hemiphragma heterophyllum Wall. | 1 | 0 | 0.19 | 0 | 0.10 | 0.44 | 0.0004 | Shrub specialist | Human valued | | |
| Pedicularis megalantha D. Don | 0.89 | 0.11 | 0.25 | 0.03 | 0.14 | 0.48 | 0.0006 | Shrubs | Human valued | Himalayan endemic | |
| Cyananthus himalaicus K. K. Shrestha | 0.94 | 0.06 | 0.24 | 0.02 | 0.13 | 0.47 | 0.0006 | Shrubs | | Nepal endemic | Threatened (Bachman et al., 2024) |
| Euphorbia stracheyi Boiss. | 0.88 | 0.12 | 0.24 | 0.03 | 0.13 | 0.46 | 0.0011 | Shrubs | Human valued | | |
| Aster sp. | 1 | 0 | 0.16 | 0 | 0.08 | 0.40 | 0.0014 | Shrub specialist | | | |
| Androsace sarmentosa Wall. | 1 | 0 | 0.16 | 0 | 0.08 | 0.40 | 0.0024 | Shrub specialist | Human valued | Himalayan endemic | |
| Sporobolus fertilis (Steud.) Clayton | 0.21 | 0.79 | 0.08 | 0.30 | 0.19 | 0.49 | 0.0029 | Open | Human valued | | |
| Epilobium sp. | 1 | 0 | 0.14 | 0 | 0.07 | 0.38 | 0.0029 | Shrub specialist | | | |
| Galium sp. | 1 | 0 | 0.14 | 0 | 0.07 | 0.38 | 0.0034 | Shrub specialist | | | |
| Delphinium himalayae Munz | 1 | 0 | 0.13 | 0 | 0.06 | 0.36 | 0.0055 | Shrub specialist | Human valued | Himalayan endemic | Threatened (Bachman et al., 2024, Ghimire et al., 2008) |
| Calamogrostis scabrescens Griseb. | 0.81 | 0.19 | 0.21 | 0.05 | 0.13 | 0.41 | 0.0130 | Shrubs | | | |
| Unknown | 1 | 0 | 0.11 | 0.00 | 0.06 | 0.33 | 0.0131 | Shrub specialist | | | |
| Arisaema jacquemontii Blume | 0.85 | 0.15 | 0.17 | 0.03 | 0.10 | 0.38 | 0.0136 | Shrubs | Human valued | | |
| Rheum acuminatum Hook. f. & Thomson | 1 | 0 | 0.11 | 0.00 | 0.06 | 0.33 | 0.0138 | Shrub specialist | Human valued | Himalayan endemic | |
| Plantago sp. | 0.15 | 0.85 | 0.03 | 0.17 | 0.10 | 0.38 | 0.0178 | Open | | | |
| Arisaema sp1 | 1 | 0 | 0.10 | 0 | 0.05 | 0.31 | 0.0270 | Shrub specialist | | | |

| Species name | Quantity 'A' | | Quantity 'B' | | | Association Statistics | *P*-value | Habitat Association | Usefulness | Endemism | Conservation Status |
|---|---|---|---|---|---|---|---|---|---|---|---|
| | Shrubs | Open | Shrubs | Open | Overall | | | | | | |
| Saxifraga hispidula D. Don | 1 | 0 | 0.08 | 0 | 0.04 | 0.28 | 0.0535 | Shrubs only - NS | | Himalayan endemic | |
| Veronica himalensis D. Don | 1 | 0 | 0.06 | 0 | 0.03 | 0.25 | 0.1134 | Shrubs only - NS | Human valued | Himalayan endemic | |
| Ligusticopsis wallichiana (DC.) Pimenov & Kljuykov | 0.86 | 0.14 | 0.10 | 0.02 | 0.06 | 0.29 | 0.1139 | NS | Human valued | | |
| Brassicaceae | 1 | 0 | 0.06 | 0.00 | 0.03 | 0.25 | 0.1179 | Shrubs only - NS | | | |
| Cyananthus lobatus Wall. ex. Benth. | 0.86 | 0.14 | 0.10 | 0.02 | 0.06 | 0.29 | 0.1189 | NS | Human valued | | |
| Cremanthodium nepalense Kitam. | 1 | 0 | 0.06 | 0.00 | 0.03 | 0.25 | 0.1218 | Shrubs only - NS | Human valued | Himalayan endemic | |
| Kobresia sp. | 0 | 1 | 0.00 | 0.06 | 0.03 | 0.25 | 0.1224 | Open only - NS | | | |
| Imperata cylindrica (L.) P. Beauv. | 0.83 | 0.17 | 0.08 | 0.02 | 0.05 | 0.26 | 0.2048 | NS | Human valued | | |
| Silene sp. | 0.83 | 0.17 | 0.08 | 0.02 | 0.05 | 0.26 | 0.2091 | NS | | | |
| Pedicularis sp. | 0.83 | 0.17 | 0.08 | 0.02 | 0.05 | 0.26 | 0.2119 | NS | | | |
| Saxifraga sp. | 1 | 0 | 0.05 | 0 | 0.02 | 0.22 | 0.2352 | Shrubs only - NS | | | |
| Impatiens harae H. Ohba & S. Akiyama | 1 | 0 | 0.05 | 0 | 0.02 | 0.22 | 0.2398 | Shrubs only - NS | | Nepal endemic | Threatened (Bachman et al., 2024) |
| Hymenidium apiolens (C.B.Clarke) Pimenov & Kljuykov | 1 | 0 | 0.05 | 0 | 0.02 | 0.22 | 0.2400 | Shrubs only - NS | | Himalayan endemic | |
| Impatiens serrata Benth. ex Hook. f. & Thomson | 1 | 0 | 0.05 | 0 | 0.02 | 0.22 | 0.2405 | Shrubs only - NS | | Himalayan endemic | |
| Cerastium fontanum Baumg. ssp. grandiflorum (Buch.-Ham. ex D.Don) H. Hara | 1 | 0 | 0.05 | 0 | 0.02 | 0.22 | 0.2422 | Shrubs only - NS | | Himalayan endemic | |
| Rhododendron lepidotum Wall. ex G. Don | 1 | 0 | 0.05 | 0 | 0.02 | 0.22 | 0.2439 | Shrubs only - NS | Human valued | | |
| Aconitum ferox Wall. ex Ser. | 1 | 0 | 0.05 | 0 | 0.02 | 0.22 | 0.2469 | Shrubs only - NS | Human valued | Himalayan endemic | Nationally threatened (Ghimire et al., 2008) |

| Species name | Quantity 'A' | | Quantity 'B' | | | Association Statistics | *P*-value | Habitat Association | Usefulness | Endemism | Conservation Status |
|---|---|---|---|---|---|---|---|---|---|---|---|
| | Shrubs | Open | Shrubs | Open | Overall | | | | | | |
| Delphinium vestitum Wall. ex Royle | 0.75 | 0.25 | 0.10 | 0.03 | 0.06 | 0.27 | 0.2653 | NS | Human valued | Himalayan endemic | |
| Persicaria sp. | 0.20 | 0.80 | 0.02 | 0.06 | 0.04 | 0.23 | 0.3544 | NS | | | |
| Rumex nepalensis Spreng. | 0.80 | 0.20 | 0.06 | 0.02 | 0.04 | 0.23 | 0.3606 | NS | Human valued | | |
| Anaphalis sp | 0.71 | 0.29 | 0.08 | 0.03 | 0.06 | 0.24 | 0.4404 | NS | Human valued | | |
| Pedicularis bifida (Buch.-Ham. ex D. Don) Pennell | 1 | 0 | 0.03 | 0 | 0.02 | 0.18 | 0.4919 | Shrubs only - NS | Human valued | Himalayan endemic | |
| Lactuca sp. | 1 | 0 | 0.03 | 0 | 0.02 | 0.18 | 0.4934 | Shrubs only - NS | | | |
| Thalictrum cultratum Wall. | 1 | 0 | 0.03 | 0 | 0.02 | 0.18 | 0.4966 | Shrubs only - NS | Human valued | | |
| Stellaria sikkimensis Hook. f. ex Edgew. & Hook. f. | 1 | 0 | 0.03 | 0 | 0.02 | 0.18 | 0.4988 | Shrubs only - NS | Human valued | Himalayan endemic | |
| Halerpestes tricuspis (Maxim.) Hnad.-Mazz. | 0.75 | 0.25 | 0.05 | 0.02 | 0.03 | 0.19 | 0.6141 | NS | Human valued | | |
| Acronema tenerum (DC.) Edgew. | 1 | 0 | 0.02 | 0 | 0.01 | 0.13 | 1 | Shrubs only - NS | | | |
| Allium wallichii Kunth. | 1 | 0 | 0.02 | 0 | 0.01 | 0.13 | 1 | Shrubs only - NS | Human valued | | Nationally threatened (Ghimire et al., 2008) |
| Anaphalis royleana DC. | 0 | 1 | 0 | 0.02 | 0.01 | 0.13 | 1 | Open only - NS | Human valued | | |
| Arisaema sp2 | 1 | 0 | 0.02 | 0 | 0.01 | 0.13 | 1 | Shrubs only - NS | | | |
| Aster himalaicus C. B. Clarke | 1 | 0 | 0.02 | 0 | 0.01 | 0.13 | 1 | Shrubs only - NS | Human valued | | |
| Cynodon sp. | 1 | 0 | 0.02 | 0 | 0.01 | 0.13 | 1 | Shrubs only - NS | | | |
| Dryopteris sp. | 1 | 0 | 0.02 | 0 | 0.01 | 0.13 | 1 | Shrubs only - NS | | | |
| Juniperus recurva Buch.-Ham. ex D. Don | 1 | 0 | 0.02 | 0 | 0.01 | 0.13 | 1 | Shrubs only - NS | Human valued | | |
| Hemipilia cucullata (L.) Y.Tang, H.Peng & T.Yukawa | 1 | 0 | 0.02 | 0 | 0.01 | 0.13 | 1 | Shrubs only - NS | | | |

| Species name | Quantity 'A' | | Quantity 'B' | | | Association Statistics | *P*-value | Habitat Association | Usefulness | Endemism | Conservation Status |
|---|---|---|---|---|---|---|---|---|---|---|---|
| | Shrubs | Open | Shrubs | Open | Overall | | | | | | |
| Parnassia nubicola Wall. ex Royle | 1 | 0 | 0.02 | 0 | 0.01 | 0.13 | 1 | Shrubs only - NS | Human valued | | Vulnerable (Ghimire et al., 2008) |
| Hymenidium benthamii (DC.) Pimenov & Kljuykov | 1 | 0 | 0.02 | 0 | 0.01 | 0.13 | 1 | Shrubs only - NS | Human valued | | |
| Rhodiola himalensis (D. Don) S. H. Fu | 1 | 0 | 0.02 | 0 | 0.01 | 0.13 | 1 | Shrubs only - NS | Human valued | | |
| Veronica sp. | 1 | 0 | 0.02 | 0 | 0.01 | 0.13 | 1 | Shrubs only - NS | | | |
| Viola biflora L. | 0.56 | 0.44 | 0.76 | 0.60 | 0.68 | 0.83 | NA | Generalists | Human valued | | |
| Taraxacum parvulum DC. | 0.31 | 0.69 | 0.41 | 0.90 | 0.66 | 0.81 | NA | Generalists | Human valued | | |
| Bupleurum candollei Wall. ex DC. | 0.43 | 0.57 | 0.52 | 0.68 | 0.60 | 0.78 | NA | Generalists | Human valued | | |
| Eriocapitella rivularis (Buch.-Ham. ex DC.) Christenh. & Byng | 0.47 | 0.53 | 0.52 | 0.59 | 0.56 | 0.75 | NA | Generalists | Human valued | | |
| Primula primulina (Spreng.) H. Hara | 0.48 | 0.52 | 0.48 | 0.51 | 0.49 | 0.70 | NA | Generalists | | | |
| Swertia angustifolia Buch.-Ham. ex D. Don | 0.68 | 0.32 | 0.62 | 0.29 | 0.45 | 0.67 | NA | Generalists | Human valued | | |
| Hedysarum sikkimense Benth. ex Baker | 0.33 | 0.67 | 0.29 | 0.59 | 0.44 | 0.66 | NA | Generalists | Human valued | | |
| Saussurea eriostemon Wall. ex C. B. Clarke | 0.67 | 0.33 | 0.57 | 0.29 | 0.43 | 0.65 | NA | Generalists | Human valued | Himalayan endemic | |
| Geranium sp. | 0.67 | 0.33 | 0.51 | 0.25 | 0.38 | 0.62 | NA | Generalists | | | |
| Potentilla sp. | 0.33 | 0.67 | 0.24 | 0.48 | 0.36 | 0.60 | NA | Generalists | | | |
| Anaphalis busua (Buch.-Ham. ex D. Don) DC. | 0.57 | 0.43 | 0.40 | 0.30 | 0.35 | 0.59 | NA | Generalists | Human valued | | |
| Bistorta macrophylla (D. Don) Sojak | 0.49 | 0.51 | 0.33 | 0.35 | 0.34 | 0.58 | NA | Generalists | Human valued | | |
| Cortia depressa (D. Don) C. Norman | 0.36 | 0.64 | 0.22 | 0.40 | 0.31 | 0.56 | NA | Generalists | Human valued | | |
| Poa sp. | 0.58 | 0.42 | 0.30 | 0.22 | 0.26 | 0.51 | NA | Generalists | | | |
| Gentiana prolata Balf.f. | 0.36 | 0.64 | 0.13 | 0.22 | 0.17 | 0.42 | NA | Generalists | Human valued | Himalayan endemic | |

| Species name | Quantity 'A' | | Quantity 'B' | | | Association Statistics | *P*-value | Habitat Association | Usefulness | Endemism | Conservation Status |
|---|---|---|---|---|---|---|---|---|---|---|---|
| | Shrubs | Open | Shrubs | Open | Overall | | | | | | |
| Oplismenus compositus (L.) P. Beauv. | 0.68 | 0.32 | 0.21 | 0.10 | 0.15 | 0.39 | NA | Generalists | | | |
| Potentilla contigua Sojak. | 0.33 | 0.67 | 0.10 | 0.19 | 0.14 | 0.38 | NA | Generalists | | | |
| Potentilla microphylla D. Don | 0.41 | 0.59 | 0.11 | 0.16 | 0.13 | 0.37 | NA | Generalists | | | |
| Leontopodium jacotianum Beauv. | 0.50 | 0.50 | 0.13 | 0.13 | 0.13 | 0.36 | NA | Generalists | Human valued | | |
| Carex sp. | 0.64 | 0.36 | 0.14 | 0.08 | 0.11 | 0.33 | NA | Generalists | | | |
| Geranium donianum Sweet | 0.64 | 0.36 | 0.14 | 0.08 | 0.11 | 0.33 | NA | Generalists | Human valued | | |
| Saxifraga kumaunensis Engl. | 0.33 | 0.67 | 0.03 | 0.06 | 0.05 | 0.22 | NA | Generalists | | | |
| Polygonaceae | 0.60 | 0.40 | 0.05 | 0.03 | 0.04 | 0.20 | NA | Generalists | | | |
| Saussurea sp. | 0.60 | 0.40 | 0.05 | 0.03 | 0.04 | 0.20 | NA | Generalists | | | |
| Anaphalis nepalensis var. monocephala (DC.) Hand.-Mazz. | 0.50 | 0.50 | 0.03 | 0.03 | 0.03 | 0.18 | NA | Generalists | Human valued | | |
| Artemisia gmelinii Weber ex Stechm. | 0.67 | 0.33 | 0.03 | 0.02 | 0.02 | 0.15 | NA | Generalists | | | |
| Bistorta amplexicaulis (D. Don) Greene | 0.67 | 0.33 | 0.03 | 0.02 | 0.02 | 0.15 | NA | Generalists | Human valued | | |
| Primula macrophylla D. Don | 0.50 | 0.50 | 0.02 | 0.02 | 0.02 | 0.13 | NA | Generalists | Human valued | | |

**Table S3: Detail list of plants recorded and their use value**

| SN | Botanical name | Family | Habit | Distribution (meters asl) | Use values | Endemism | Conservation status (Reference) |
|---|---|---|---|---|---|---|---|
| **1** | *Aconitum ferox* Wall. ex Ser. | Ranunculaceae | Herb | 2100-4700 | Medicine, Social, Commercial | Himalayan endemic | Nationally threatened (Ghimire et al. 2008) |
| **2** | *Acronema tenerum* (DC.) Edgew. | Apiaceae | Herb | 2100-4500 | | | |
| **3** | *Allium wallichii* Kunth. | Amaryllidaceae | Herb | 2100-4800 | Medicine, Food | | Nationally threatened (Ghimire et al. 2008) |
| **4** | *Anaphalis busua* (Buch.-Ham. ex D. Don) DC. | Asteraceae | Herb | 700-3500 | Medicine, Social | | |
| **5** | *Anaphalis nepalensis* (Spreng.) Hand.-Mazz. | Asteraceae | Herb | 2700-4700 | Medicine, Social, Food (leaves) | | |
| **6** | *Anaphalis nepalensis* var. *monocephala* (DC.) Hand.-Mazz. | Asteraceae | Herb | 3500-5300 | Medicine | | |
| **7** | *Anaphalis royleana* DC. | Asteraceae | Herb | 2600-4300 | Medicine | | |
| **8** | Anaphalis sp | Asteraceae | Herb | | | | |
| **9** | *Androsace sarmentosa* Wall. | Primulaceae | Herb | 2500-4000 | Medicine | Himalayan endemic | |
| **10** | *Arisaema jacquemontii* Blume | Araceae | Herb | 2400-4500 | Medicine, Food, Social | | |
| **11** | *Arisaema* sp1 | Araceae | Herb | | | | |
| **12** | *Arisaema* sp2 | Araceae | Herb | | | | |
| **13** | *Artemisia gmelinii* Weber ex Stechm. | Asteraceae | Shrub | 1300-4300 | | | |
| **14** | *Aster himalaicus* C. B. Clarke | Asteraceae | Herb | 3500-5200 | Medicine | | |
| **15** | *Aster* sp. | Asteraceae | Herb | | | | |
| **16** | *Bistorta amplexicaulis* (D. Don) Greene | Polygonaceae | Herb | 1400-4200 | Medicine | | |
| **17** | *Bistorta macrophylla* (D. Don) Sojak | Polygonaceae | Herb | 2300-4700 | Medicine, Food | | |
| **18** | Brassicaceae | Brassicaceae | Herb | | | | |
| **19** | *Bupleurum candollei* Wall. ex DC. | Apiaceae | Herb | 1500-4900 | Fodder (but tender leaves poisonous to cattle) | | |
| **20** | *Calamagrostis pseudophragmites* (Haller f.) Koeler | Poaceae | Herb | 1500-4600 | | | |
| **21** | Calamogrostis scabrescens Griseb. | Poaceae | Herb | 2400-4700 | | | |
| **22** | *Caltha palustris* L. | Ranunculaceae | Herb | 2400-4600 | Medicine (but roots are poisonous) | | |
| **23** | *Carex* sp. | Cyperaceae | Herb | | | | |
| **24** | *Cerastium fontanum* Baumg. ssp. grandiflorum (Buch.-Ham. ex D.Don) H. Hara | Caryophyllaceae | Herb | 1900-3800 | | Himalayan endemic | |

| SN | Botanical name | Family | Habit | Distribution (meters asl) | Use values | Endemism | Conservation status (Reference) |
|---|---|---|---|---|---|---|---|
| **25** | *Cortia depressa* (D. Don) C. Norman | Apiaceae | Herb | 2900-5200 | Medicine, Food, Food additives | | |
| **26** | *Corydalis juncea* Wall. | Papaveraceae | Herb | 2500-5100 | Medicine | Himalayan endemic | |
| **27** | *Cremanthodium nepalense* Kitam. | Asteraceae | Herb | 2900-5200 | Medicine | Himalayan endemic | |
| **28** | *Cyananthus himalaicus* K. K. Shrestha | Campanulaceae | Herb | 3000-4800 | | Endemic to Nepal | Threatened (Bachman et al., 2024) |
| **29** | *Cyananthus lobatus* Wall. ex. Benth. | Campanulaceae | Herb | 3300-4700 | Medicine, Social | | |
| **30** | *Cynodon* sp. | Poaceae | Herb | | | | |
| **31** | *Cyperus* sp. | Cyperaceae | Herb | | | | |
| **32** | *Delphinium himalayae* Munz | Ranunculaceae | Herb | 2000-4600 | Medicine, Commercial | Himalayan endemic | Threatened (Bachman et al., 2024, Ghimire et al., 2008) |
| **33** | *Delphinium vestitum* Wall. ex Royle | Ranunculaceae | Herb | 2700-4700 | Medicine, Poisonous (for dogs) | Himalayan endemic | |
| **34** | *Dryopteris* sp. | Dryopteridaceae | Fern | | | | |
| **35** | *Epilobium* sp. | Onagraceae | Herb | | | | |
| **36** | Eriocapitella rivularis (Buch.-Ham. ex DC.) Christenh. & Byng | Ranunculaceae | Herb | 1600-4400 | Medicine, Food, Poison (Insecticidal, whole plant) | | |
| **37** | *Euphorbia stracheyi* Boiss. | Euphorbiaceae | Herb | 2100-5000 | Medicine | | |
| **38** | *Galium* sp. | Rubiaceae | Herb | 2700-3600 | | | |
| **39** | *Gentiana depressa* D. Don | Gentianaceae | Herb | 2900-4300 | Medicine | | |
| **40** | *Gentiana prolata* Balf.f. | Gentianaceae | Herb | 3100-5500 | Medicine | Himalayan endemic | |
| **41** | *Geranium donianum* Sweet | Geraniaceae | Herb | 3200-4800 | Medicine | | |
| **42** | *Geranium* sp. | Geraniaceae | Herb | | | | |
| **43** | Halerpestes tricuspis (Maxim.) Hnad.-Mazz. | Ranunculaceae | Herb | 2600-4600 | Medicine | | |
| **44** | *Hedysarum sikkimense* Benth. ex Baker | Fabaceae | Herb | 3100-4700 | Medicine | | |
| **45** | *Hemiphragma heterophyllum* Wall. | Plantaginaceae | Herb | 1300-3700 | Medicine, Food | | |
| **46** | *Hemipilia cucullata* (L.) Y.Tang, H.Peng & T.Yukawa | Orchidaceae | Herb | 3700-5000 | | | |
| **47** | *Hymenidium apiolens (C.B.Clarke) Pimenov & Kljuykov* | Apiaceae | Herb | 3300-4500 | | Himalayan endemic | |

| SN | Botanical name | Family | Habit | Distribution (meters asl) | Use values | Endemism | Conservation status (Reference) |
|---|---|---|---|---|---|---|---|
| **48** | *Hymenidium benthamii (DC.) Pimenov & Kljuykov* | Apiaceae | Herb | 2200-4700 | Medicine | | |
| **49** | *Impatiens harae* H. Ohba & S. Akiyama | Balsaminaceae | Herb | 1900-3700 | | Endemic to Nepal | Threatened (Bachman et al., 2024) |
| **50** | *Impatiens serrata* Benth. ex Hook. f. & Thomson | Balsaminaceae | Herb | 2000-3600 | | Himalayan endemic | |
| **51** | *Impatiens* sp. | Balsaminaceae | Herb | | | | |
| **52** | *Imperata cylindrica* (L.) P. Beauv. | Poaceae | Herb | 100-2400 | Medicine, Fodder, Materials | | |
| **53** | *Juniperus recurva* Buch.-Ham. ex D. Don | Cupressaceae | Shrub | 1900-4600 | Medicine, Social, Fuel, Commercial | | |
| **54** | *Kobresia* sp. | Cyperaceae | Herb | | | | |
| **55** | *Lactuca* sp. | Asteraceae | Herb | | | | |
| **56** | *Leontopodium jacotianum* Beauv. | Asteraceae | Herb | 2700-4900 | Medicine, Social | | |
| **57** | *Ligusticopsis wallichiana* (DC.) Pimenov & Kljuykov | Apiaceae | Herb | 1400-5000 | Medicine, Food, Food additives, Commercial | | |
| **58** | *Oplismenus compositus* (L.) P. Beauv. | Poaceae | Herb | 100-2800 | | | |
| **59** | *Parnassia nubicola* Wall. ex Royle | Celastraceae | Herb | 2100-4600 | Medicine, Commercial | | Vulnerable (Ghimire et al., 2008) |
| **60** | *Pedicularis bifida* (Buch.-Ham. ex D. Don) Pennell | Orobanchaceae | Herb | 1000-3600 | Medicinal | Himalayan endemic | |
| **61** | *Pedicularis megalantha* D. Don | Orobanchaceae | Herb | 2500-4300 | Medicine | Himalayan endemic | |
| **62** | *Pedicularis* sp. | Scrophulariaceae | Herb | | | | |
| **63** | *Persicaria* sp. | Polygonaceae | | | | | |
| **64** | *Plantago* sp. | Plantaginaceae | Herb | | | | |
| **65** | *Poa* sp. | Poaceae | | | | | |
| **66** | Polygonaceae | Polygonaceae | Herb | | | | |
| **67** | *Potentilla contigua* Sojak. | Rosaceae | Herb | 3500-4700 | | | |
| **68** | *Potentilla daltoniana* (J. Gay) Mabb. | Rosaceae | Herb | 2600-4000 | Medicine, Food | | |
| **69** | *Potentilla microphylla* D. Don | Rosaceae | Herb | 3400-5200 | | | |
| **70** | *Potentilla* sp. | Rosaceae | Herb | | | | |
| **71** | *Primula macrophylla* D. Don | Primulaceae | Herb | 3300-5800 | Medicine | | |
| **72** | *Primula primulina* (Spreng.) H. Hara | Primulaceae | Herb | 2100-5000 | | | |
| **73** | *Rheum acuminatum* Hook. f. & Thomson | Polygonaceae | Herb | 2800-4600 | Medicine, food, material | Himalayan endemic | |
| **74** | *Rhodiola himalensis* (D. Don) S. H. Fu | Crassulaceae | Herb | 3300-4800 | Medicine | | |

| SN | Botanical name | Family | Habit | Distribution (meters asl) | Use values | Endemism | Conservation status (Reference) |
|---|---|---|---|---|---|---|---|
| **75** | *Rhododendron lepidotum* Wall. ex G. Don | Ericaceae | Shrub | 2100-4700 | Medicine, Social, Food, Spray | | |
| **76** | *Rumex nepalensis* Spreng. | Polygonaceae | Herb | 200-4200 | Medicine, Food | | |
| **77** | *Saussurea eriostemon* Wall. ex C. B. Clarke | Asteraceae | Herb | 3200-4900 | Medicine, Social, Materials | Himalayan endemic | |
| **78** | *Saussurea* sp. | Asteraceae | Herb | | | | |
| **79** | *Saxifraga diversifolia* Wall. ex Ser. | Saxifragaceae | Herb | 2400-4800 | Medicine | | |
| **80** | *Saxifraga hispidula* D. Don | Saxifragaceae | Herb | 3000-4500 | | Himalayan endemic | |
| **81** | *Saxifraga kumaunensis* Engl. | Saxifragaceae | Herb | 3300-4800 | | | |
| **82** | *Saxifraga* sp. | Saxifragaceae | Herb | | | | |
| **83** | *Silene* sp. | Caryophyllaceae | Herb | | | | |
| **84** | *Sporobolus fertilis* (Steud.) Clayton | Poaceae | Herb | 300-3800 | Fodder, Materials | | |
| **85** | *Stellaria patens* D. Don | Caryophyllaceae | Herb | 1300-4200 | | Himalayan endemic | |
| **86** | *Stellaria sikkimensis* Hook. f. ex Edgew. & Hook. f. | Caryophyllaceae | Herb | 2000-3800 | Medicine | Himalayan endemic | |
| **87** | *Swertia angustifolia Buch.-Ham. ex D. Don* | Gentianaceae | Herb | 600-3800 | Medicine, commercial. | | |
| **88** | *Taraxacum parvulum* DC. | Asteraceae | Herb | 800-3000 | Medicine, Food | | |
| **89** | *Tetrataenium wallichii* (DC.) Manden. | Apiaceae | Herb | 2500-4300 | Medicinal | Himalayan endemic | Threatened (Bachman et al., 2024) |
| **90** | *Thalictrum cultratum* Wall. | Ranunculaceae | Herb | 1400-4500 | Medicine | | |
| **91** | *Themeda triandra* Forssk. | Poaceae | Herb | 1700-4200 | Materials | | |
| **92** | Unknown | ? | | | | | |
| **93** | *Unknown2* | ? | Herb | | | | |
| **94** | *Valeriana hardwickii* Wall. | Caprifoliaceae | Herb | 1200-4200 | Medicine, social, food. | | |
| **95** | *Veronica himalensis* D. Don | Plantaginaceae | Herb | 2600-5000 | Medicine | Himalayan endemic | |
| **96** | *Veronica* sp. | Plantaginaceae | Herb | | | | |
| **97** | *Viola biflora* L. | Violaceae | Herb | 2100-4600 | Medicine, Materials | | |
